# Multimodal AI/ML for discovering novel biomarkers and predicting disease using multi-omics profiles of patients with cardiovascular diseases

**DOI:** 10.1101/2024.08.07.607041

**Authors:** William DeGroat, Habiba Abdelhalim, Elizabeth Peker, Neev Sheth, Rishabh Narayanan, Saman Zeeshan, Bruce T. Liang, Zeeshan Ahmed

## Abstract

Cardiovascular diseases (CVDs) are multifactorial diseases, requiring personalized assessment and treatment. The advancements in multi-omics technologies, namely RNA-seq and whole genome sequencing, have offered translational researchers a comprehensive view of the human genome; utilizing this data, we can reveal novel biomarkers and segment patient populations based on personalized risk factors. Limitations in these technologies in failing to capture disease complexity can be accounted for by using an integrated approach, characterizing variants alongside expression related to emerging phenotypes. Designed and implemented data analytics methodology is based on a nexus of orthodox bioinformatics, classical statistics, and multimodal artificial intelligence and machine learning techniques. Our approach has the potential to reveal the intricate mechanisms of CVD that can facilitate patient-specific disease risk and response profiling. We sourced transcriptomic expression and variants from CVD and control subjects. By integrating these multi-omics datasets with clinical demographics, we generated patient-specific profiles. Utilizing a robust feature selection approach, we reported a signature of 27 transcripts and variants efficient at predicting CVD. Here, differential expression analysis and minimum redundancy maximum relevance feature selection elucidated biomarkers explanatory of the disease phenotype. We used Combination Annotation Dependent Depletion and allele frequencies to identify variants with pathogenic characteristics in CVD patients. Classification models trained on this signature demonstrated high-accuracy predictions for CVDs. Overall, we observed an XGBoost model hyperparameterized using Bayesian optimization perform the best (AUC 1.0). Using SHapley Additive exPlanations, we compiled risk assessments for patients capable of further contextualizing these predictions in a clinical setting. We discovered a 27-component signature explanatory of phenotypic differences in CVD patients and healthy controls using a feature selection approach prioritizing both biological relevance and efficiency in machine learning. Literature review revealed previous CVD associations in a majority of these diagnostic biomarkers. Classification models trained on this signature were able to predict CVD in patients with high accuracy. Here, we propose a framework generalizable to other diseases and disorders.

## 1. Introduction

Cardiovascular diseases (CVDs) are recognized as the primary cause of mortality among men and women in the United States [1, 2]. Given the complex nature, risk factors, inherent genetic makeup, and trajectory of CVD, personalized management is essential for effective treatment [2]. Advancements in genomics and bioinformatics have significantly enhanced our understanding of the intricate origins of CVDs [3, 4]. Gaining insights into disease implications by utilizing transcriptomic expression and variant profiles holds the promise of revolutionizing diagnostic capabilities, treatment strategies, and prognostic assessments across various CVDs including but not limited to heart failure (HF) and arial fibrillation (AF) [5, 6]. These advancements stem from next-generation sequencing (NGS) technologies, which have facilitated the identification of novel heritable links and the exploration of genetic diversity among patients [7]. Gene expression analysis through RNA-seq data has aided in uncovering disease associated biomarkers and categorizing patient groups according to their risk profiles [8]. Analyzing RNA-seq data for differential expression allows for the exploration of genome-wide biological disparities, leading to enriched functional pathways and gene ontologies [9, 10].

RNA-seq data provides valuable biological insights into gene expression, RNA processing, and molecular pathways underlying disease states [11, 12]. While gene expression analysis allows for enhancements in diagnostic capabilities and precise treatment plans, multiple studies have established that RNA-seq provides limited coverage of non-coding regions, and that transcriptomics cannot detect genomic variants [11, 12, 13]. The onset of multifactorial diseases is shaped by an interplay of environmental and genetic factors, affecting various biological processes such as gene regulation [14]. Previous studies utilizing whole genome and exome sequencing (WGS/WES) have demonstrated their efficacy in accurately revealing the effects of non-coding variants on CVDs [15, 16] and other complex diseases [17], as well as in capturing all genetic variation. Thus, providing comprehensive information about an individual’s entire genome [16, 17]. Although sequencing technology aids in identifying genetic variations linked to diseases, accurately linking specific genomic variations to disease phenotypes remains challenging [3, 18]. Deciphering the pathogenic and biological function of genes may require additional information beyond what one type of data can offer [18]. Data integration is vital in managing the escalating volume of data and obtaining comprehensive interdisciplinary insights into extensive genomic datasets [19]. Additionally, due to the heterogenous nature of genomic, transcriptomic, and clinical data, there is a lack of standardization creating a persistent limitation in data integration [18]. These challenges are being addressed with the integration of precision medicine and artificial intelligence (AI)/machine learning (ML) approaches where phenotypic, clinical, transcriptomic, and genomic data can be subjected to classification and selection to facilitate the identification of high-risk patients [18, 20]. Utilizing cutting edge AI/ML technology can aid in the analysis and interpretation of gene expression and variant data, providing more accurate diagnosis and improving our understanding of the mechanisms behind complex diseases including but not limited to CVDs [21, 22], lupus [23], and colon cancer [24].

Previously, we have performed traditional bioinformatic analyses including an in-depth gene expression and enrichment analysis of RNA-seq data from patients with mostly HF and other CVDs. We identified differentially expressed genes (DEGs) that are well-documented to be associated with CVDs and other enriched pathways [25]. However, we were unable to detect any CVD drivers utilizing RNA-seq data. To address this limitation, we employed an integrative, multi-omics approach of gene expression, disease-causing gene variants and associated phenotypes among CVD populations [6]. In this study we combined specific mutations for the DEGs we had previously reported allowing for a better understanding of CVD progression [6]. Extending our research and expanding beyond orthodox bioinformatic techniques, we implemented AI/ML techniques on RNA-seq driven gene expression data to study biomarkers associated with HF, AF, and other CVDs [26]. Our AI/ML analysis supported our initial gene expression study as we were able to identify common genes that have a high impact on CVD diagnosis [25, 26]. Additionally, this AI/ML framework aided in establishing *Hygieia*, a portable pipeline which integrates genomics and healthcare data to explore genes linked to specific disorders and predict disease [27]. While we were able to predict CVDs with high accuracy using this methodology, we were only focused on CVD driver genes with genetic alterations that can culminate in CVD [26]. We overcame this challenge by using whole transcriptome-based gene expression data and further enhanced our AI/ML model to adapt a novel nexus of algorithms to predict CVDs based on crucial transcriptomic biomarkers [28]. We utilized this approach and proposed *IntelliGenes*, a novel AI/ML pipeline for the identification of novel biomarkers and predict disease in individuals [29].

In this study, we leverage our previous work and present a new AI/ML approach that uses multi-omics data integrating RNA-seq driven gene expression, whole genome based single nucleotide variant (SNVs), and demographic and clinical data (Figure 1). Novel biomarkers based on differential expression associated with CVDs were investigated for pathogenic SNVs to identify variations within the genes and their regulatory elements. A clinically integrated genomic and transcriptomic (CIGT) dataset was leveraged by two classifiers and three ML algorithms to accurately predict CVDs. Through the identification of genetic biomarkers and their relative SNPs, we have highlighted potential indicators for the early detection of CVDs. These biomarkers aid in identifying individuals at risk pre-diagnosis, enabling prompt intervention, and enhancing patient outcomes. With its implementation in healthcare, our predictive model can identify patients at risk of CVDs and may be extrapolated to perform other single disease predictions.

**Figure 1.**
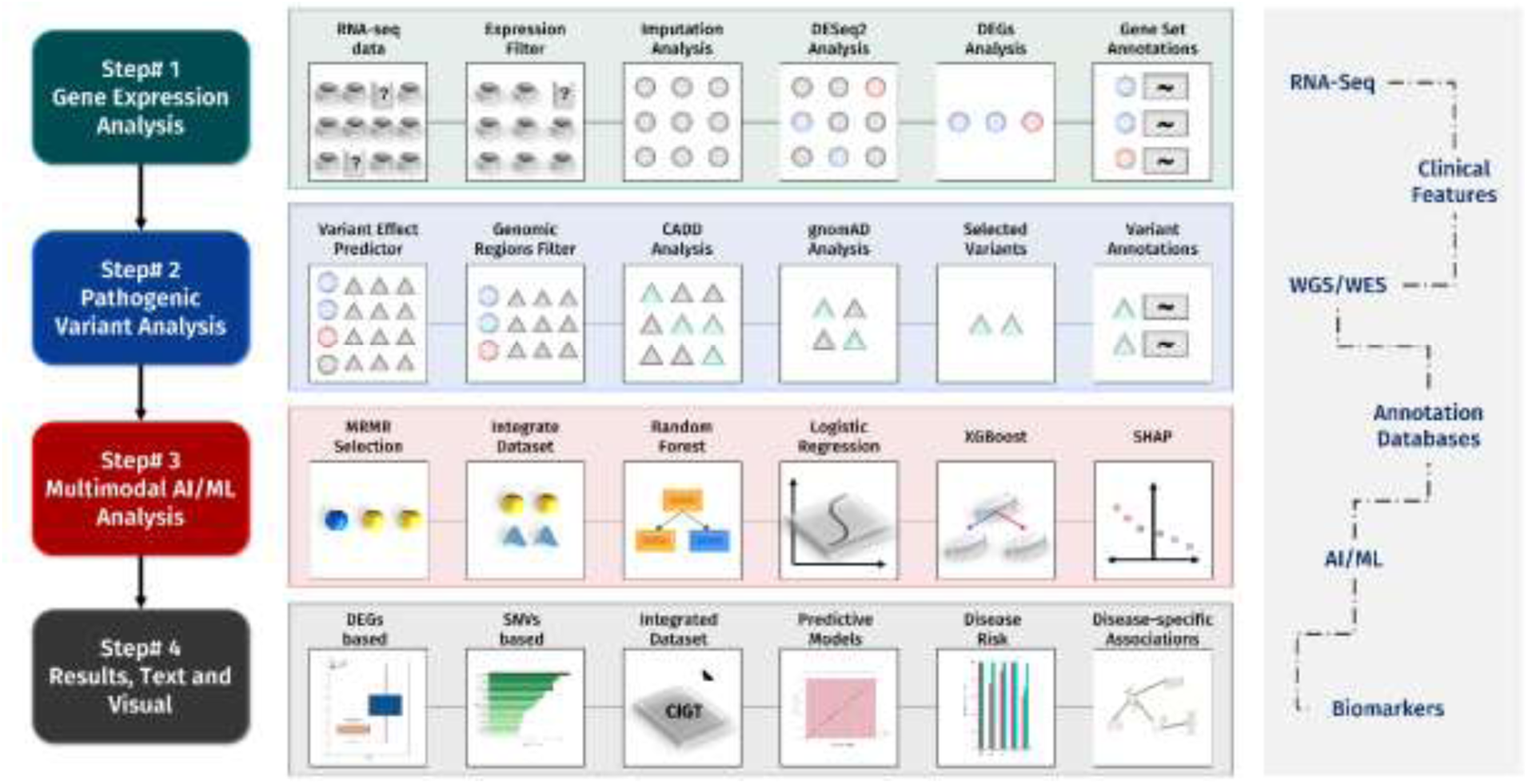
Study design and workflow. This figure represents a summary of our study design: I) Transcriptomic expression, II) Pathogenic Variant, III) Multimodal Machine Learning, and IV) Results. Various inputs and their implementation are also included (RNA-sequencing, Clinical Records, Whole Genome Sequencing, Annotation databases and Biomarkers).

## 2. Methods

Overall methodology is into three steps (Figure 2), which include I) transcriptomic/gene expression analysis, II) pathogenic variant analysis, and III) Multimodal AI/ML analysis.

**Figure 2.**
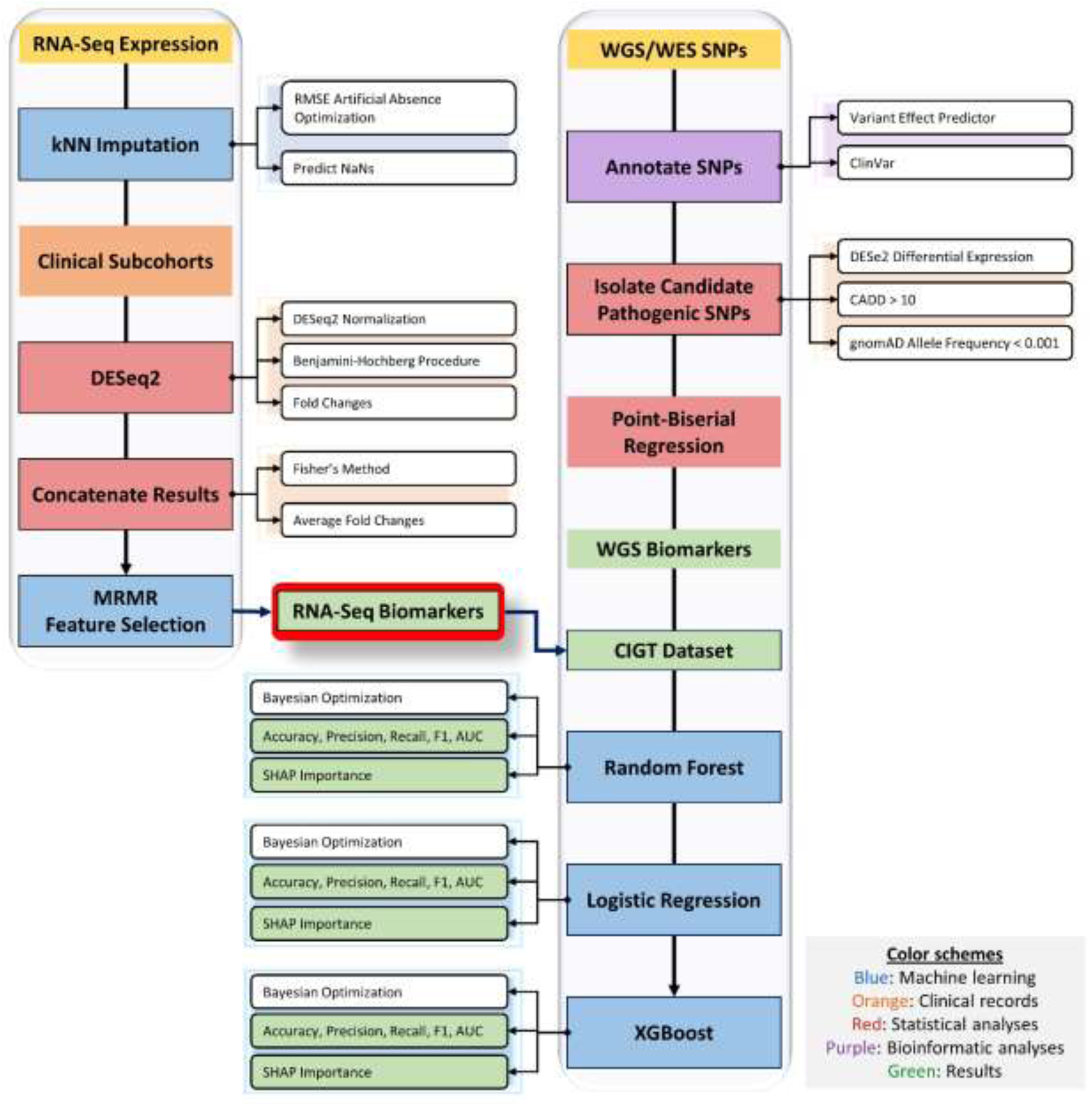
Methodology. This figure presents the k-Nearest Neighbors (k-NN) imputation to address missing values present in our RNA-seq expression data. DESeq2 was utilized for normalization and gene differential expression on four clinical sub cohorts to reduce the effect of confounding variables. Next, minimum redundancy – maximum relevance (MRMR) was performed to identify biomarkers proficient in predicting CVDs. Simultaneously, significant single nucleotide variants (SNVs) were annotated, and their pathogenicity determined for downstream analysis. Utilizing the clinically integrated transcriptomics and genomics dataset (CIGT) of significant biomarkers and their variants, machine learning algorithms (Random Forest, Logistic Regression, and Xtreme Gradient Boosting) to predict CVDs. Boxes highlighted in yellow refer to input data, blue refers to machine learning approaches, orange highlights clinical records, red refers to statistical analyses, while purple refers to bioinformatic analyses, and green highlights results.

### Transcriptomic/Gene Expression Analysis

Previously, we performed RNA-seq on a 71-participant cohort of CVD patients and health controls [6, 25]. Samples were collected from the individual’s peripheral blood mononuclear cells (PBMCs). All procedures performed in studies involving human participants were in accordance with the ethical standards of the institution and with the 1964 Helsinki Declaration and its later amendments or comparable ethical standards. All human samples were used in accordance with relevant guidelines and regulations, and all experimental protocols were approved by the Institutional Review Board. This cohort had 61 CVD patients: 40 male and 21 female individuals aged between 45 and 92, from diverse ethnic groups (56 non-Hispanic, four Hispanic, and one declined to answer) and self-described race (42 Whites, seven Blacks or African Americans, one Asian and 11 of unknown race). Ten control patients rounded out the cohort: five males and five females, (out of which three were self-described Hispanics and seven non-Hispanic; nine were White race and one unknown race) aged between 28 and 78 with no clinical manifestation of CVD. From our 71-participant cohort, we extracted transcriptomic expression for each individual. Counts and TPM values were retrieved from our RNA-seq. TPM values guided preprocessing. Transcripts with a median TPM below 0.5 across participants were removed from our dataset, and transcripts without significant expression (TPM > 1) in at least one patient were also removed. Additionally, non-ubiquitous transcripts, or transcripts expressed in less than 80% of the cohort’s participants’ RNA-seq, were excluded from downstream analysis. The filtered transcripts were configured into a CIGT-formatted dataset based on raw count values; this dataset served as the basis of our series of analyses.

We applied k-nearest neighbors (k-NN) imputation paired with artificial missingness to predict missing count values within our CIGT-formatted dataset (e.g., NaN). Here, we replaced 10% of the known portion of our dataset with missing values, maintaining the true values for comparison. We simulated imputations using distinct ‘n_neighbors’ values, ranging from 1 to 20. Optimizing this parameter assists in reducing noise in high-dimensional datasets, fine-tuning the imputer’s sensitivity; essentially, this parameter tells the algorithm the number of most similar data points to consult when estimating a missing value. With each simulation, we calculated the root mean squared error (RMSE) by comparing the predicted values with the previously withheld portion of the dataset; RMSE was minimized in our choice of optimal ‘n_neighbors’ values. We filled in missing values using optimal ‘n_neighbors’ and reintegrated the artificially excluded values into our dataset. We conducted dataset normalization using DESeq2’s median of ratios on our continuous imputed dataset [30, 31]. This normalization method, which utilizes RNA-seq counts, adjusts for cross-sample comparability better than TPM values [32]. We calculated the coefficient of variation and intraclass correlation for our normalized dataset. Because median of ratios is a dataset-dependent normalization algorithm, sub-setted matrices from our expression data must be renormalized from raw counts prior to further analysis.

Next, we performed differential expression analysis using DESeq2. To minimize the effects of confounders, the cohort’s CVD patients and control participants were stratified into subcohorts based on their demographic features. We assigned groups using the individual’s sex, racial background, and age. Four groups were created: white males ages 45 - 64, white males ages 65+, white females ages 45 - 64, and white females 65 plus. Differential expression using DESeq2 was then performed independently on each subcohort, employing a negative binomial distribution-based model to locate DEGs. [30]. DE on each subcohort yielded separate p-values and log fold changes (LFC) per feature, respectively indicating its probability of being differentially expressed and the directionality of expression. A positive LFC value indicates up regulation (i.e. over-expression), and negative indicates down regulation (i.e. under-expression). The results from our four subcohorts are merged: p-values were combined with Fisher’s method and LFCs were averaged. Only DEGs with an adjusted p-value less than 0.05 were considered to be significant and included in downstream analysis.

To determine biologically relevant DEGs use in ML single-disease predictions [33, 34], and to select genes explanatory of a generalized CVD phenotype, we utilized minimum redundancy -maximum relevance (MRMR). In MRMR, an arbitrary ‘k’ parameter is chosen indicating the number of biomarkers returned. This is the best set of ‘k’ size for explaining the difference between patients and controls. Here, MRMR minimizes the effects of co-expressed DEGs, and other patterns of redundant information contained in expression datasets. This example is preferable to other ML selectors with arbitrary cutoffs, such as recursive feature elimination models; in such models, highly correlated biomarkers might aggregate atop rankings, leaving classifiers with less useful or more redundant information to make predictions. Here, we performed MRMR exclusively on the DEGs in our training dataset, with ‘k’ set as 10. Using Gene Set Enrichment Analysis (GSEA), we examined Gene Ontology (GO) and Human Phenotype Ontology (HPO) enrichment in our 10 MRMR-selected biomarkers [35, 36, 37]. GO was utilized to investigate the biological processes, cellular components, and molecular functions of our biomarkers. HPO assisted in searching for disease implications.

### Pathogenic Variant Analysis

WGS from our 71-participant cohort was investigated alongside RNA-seq. We processed WGS-derived single nucleotide polymorphisms (SNPs) from each patient into VCF format. SNPs with low quality scores (< 50) within the VCF files were discarded. Using the Ensembl Variant Effect Predictor (VEP), we annotated these genomes with ClinVar, Combined Annotation Dependent Depletion (CADD), and Genome Aggregation Database (gnomAD). Here, we exclusively examined the WGS datasets of CVD patients in our training cohort for these SNPs. Only SNPs associated with our MRMR-selected DEGs and their known regulatory elements (i.e., promoters and enhancers), sourced from GeneHancer [38], were included in downstream analyses. This methodology allowed us to focus on genomic regions we had previously been able to implicate with CVDs. By excluding SNPs outside these key areas, our study minimizes confounding data, thereby improving the likelihood of identifying significant genetic contributors to CVD.

Next, we utilized CADD to measure the deleteriousness of SNPs in our regions of interest. CADD is a computational tool for understanding the pathogenicity of SNPs [39]. CADD utilizes machine learning to integrate multiple genomic annotations, predicting the perniciousness of genetic variants. This tool leverages data from diverse sources, including evolutionary conservation and functional annotations, to generate a comprehensive score that assesses variant impact. We used a CADD Phred score greater than 10, or the top 10% of harmful SNPs, as our threshold. gnomAD was utilized to discover rare variants within this highly deleterious set. gnomAD is a comprehensive public resource that aggregates exome and genome sequencing data to provide insights into genetic variants across diverse populations. It offers critical information on the frequency and potential impact of genetic mutations. gnomAD provided allele frequencies for SNPs; those with an allele frequency < 0.1% were included in our analyses [40]. We searched for the presence of SNPs from our training dataset in the rest of our 71-patient cohort. The presence/absence of these SNPs was detailed in a binary matrix in CIGT format. This matrix was merged with our RNA-seq counts matrix by a common ID identifying each patient. To examine if our selected SNPs showed any direct linear relation with transcriptomic expression, we performed a point-biserial correlation.

### Multimodal AI/ML Analysis

Our selected DEGs and SNPs were integrated into an AI/ML-ready CIGT-formatted dataset and used with an 80/20 train-test split by three ML classifiers to predict CVD risk. Using Bayesian optimization, we found optimal hyperparameters for a random forest (RF), Xtreme Gradient Boosting (XGBoost), and logistic regression (LR) models. For RF, ‘max_depth’, ‘min_samples_split’, ‘min_samples_leaf’, ‘n_estimators’, ‘max_features’, ‘bootstrap’, ‘criterion’, ‘max_leaf_nodes’, and ‘max_samples’ were optimized. Our XGBoost classifier was optimized for ‘max_depth’, ‘min_child_weight’, ‘gamma’, ‘subsample’, ‘colsample_bytree’, ‘scale_pos_weight’, ‘n_estimators’, and ‘learning_rate’. The parameters ‘C’, ‘penalty’, ‘solver’, and ‘max_iter’ were included in the Bayesian optimization for LR. Optimizing these hyperparameters assists in maximizing the performance of each classifier on the dataset. Bayesian optimization is a specific sequential optimization technique enabling faster convergence in parameter over typical brute force algorithms such as Grid Search. RF and XGBoost, two tree-based models, were chosen as they have previously proven powerful in single-disease prediction [28]. LR, a linear model, was chosen for comparison. These classifiers performed patient single-disease prediction on our testing dataset. Metrics detailing the classifier’s performance were computed: accuracy, AUC, probabilities, sensitivity, specificity, F1, and Brier score. To investigate the importance and directionality of each feature in predicting the CVD phenotype for each ML model, SHapley Additive exPlanations (SHAP) were computed for each feature. SHAP scores offer insights into patient-specific CVD manifestations using a game theoretic approach [41]. We combined the SHAP profiles with prediction probabilities to investigate which biomarkers were the most important contributors to each patient’s CVD prediction. Additionally, we extensively reviewed literature search to examine which of these biomarkers have previously been implicated in manifestations of CVD.

## 3. Results

We designed comparable, RNA-seq data driven patient-specific expression profiles efficient in DE and ML analyses [29]. We found 56,681 transcripts relevant to the DEGs in our cohort. Initial filtration revealed only 748 transcripts for follow up differential expression analysis. Missing expression values in individual patients’ profiles necessitated imputation before DEA. Using a robust, empirical approach to parameterizing an imputer, we corrected dataset absence. Artificial missingness facilitated RMSE calculations across various simulations of ‘n_neighbors’. K-NN Imputation was performed using our optimal ‘n_neighbors’ of 11, having the lowest RMSE across simulations. With a completed dataset, DESeq2 normalization was performed for improved cross-sample comparability. Utilizing a demographic-based segmentation approach, we performed DEA to investigate potential CVD-associated transcripts in four subcohorts, minimizing noise from confounding variables. Subcohort-specific results from DEA were then merged, detecting 28 DEGs, detailed in Table 1, across our cohort. LFCs characterized each DEG’s direction of regulation in CVD patients. Here, we demonstrate 20 upregulated DEGs and 8 downregulated DEGs.

**Table 1.** Analysis and Associations of differentially expressed genes. This table includes differentially expressed genes, their p-values to determine significance, log fold change (LFC) to determine regulation (up/down), direct cardiovascular and non-cardiovascular disease (CVD) associations based on current and extensive literature review. Genes highlighted in green correspond to a direct CVD association, blue represents a direct CVD and non-CVD relationship, and yellow corresponds to a non-CVD association.

| # | Gene | p-value | Log fold change (LFC) | Regulation (Up/Down) | CVD association | Non-CVD Association |
| --- | --- | --- | --- | --- | --- | --- |
| 1 | HBM | 0.00079763 | 2.646956203 | Up | Hypertrophic cardiomyopathy | No Association |
| 2 | GUK1 | 0.007690665 | 2.046060722 | Up | Coronary heart disease, stroke, ischemic stroke, hypertrophic cardiomyopathy | No Association |
| 3 | HBA1 | 0.003253392 | 2.088482737 | Up | Ischemic heart disease, ischemic stroke, heart failure, coronary artery disease | Non-alcoholic fatty liver disease, chronic kidney disease |
| 4 | GPX1 | 0.007653606 | 1.580476992 | Up | Cardiomyopathy | Kashin-beck disease, renal papillary cell carcinoma, ovarian serous cystadenocarcinoma, pancreatic adenocarcinoma, skin melanoma, thyroid cancer, endometrial cancer |
| 5 | SELENBP1 | 0.022601729 | 2.105152398 | Up | Myocardial infarction, cardiac arrest, myocardial hypoxia, myocardial ischemia, acute coronary syndrome | Sideroblastic anemia, breast cancer, prostate cancer, renal cell cancer, colorectal cancer, lung adenocarcinomas |
| 6 | LGALS3 | 0.042676564 | 1.704069001 | Up | Heart failure, ischemic stroke, coronary heart disease | Colorectal cancer, pancreatic cancer |
| 7 | ND1 | 0.049145898 | 1.950520657 | Up | Maternally inherited coronary heart disease, cardiomyopathy | Mitochondrial encephalomyopathy, lactic acidosis |
| 8 | ITGB2 | 3.27E-05 | -1.914719723 | Down | Coronary heart disease, acute myocardial infarction | Non-small cell lung cancer, leukocyte adhesion deficiency type-1, low-grade glioma, ovarian cancer, rheumatoid arthritis, osteoarthritis, Hirschsprung's disease-associated enterocolitis |
| 9 | ACTB | 0.031950547 | -2.269915072 | Down | Coronary heart disease, stroke, ischemic stroke, hypertrophic cardiomyopathy | Pleiotropic developmental disorder, keratoconus, dystonia-deafness, late-onset Parkinson's disease |

We implemented MRMR feature selection to capture the phenotypic profile of CVD in 10 DEGs: *ITGB2*, *CD37*, *RPL36AP37*, *PSAP*, *ACTB*, *SELL*, *NCF2*, *HBA1*, *ICAM3*, and *BBLN*. This excludes transcriptomic features with non-informative and redundant information to ML classifiers (e.g., co-expression). As demonstrated, these 10 DEGs successfully explain the differences between CVD patients and healthy controls. Figures 3A and 3B detail each DEG’s LFC and adjusted p-value. Additionally, we examined GO and HPO enrichment across the transcript set to examine their implicated pathways and clinical relevance (Figure 3C). Using Fisher’s exact test, we conclude that downregulated DEGs are enriched in our MRMR-selected DEGs, with a p-value of 0.022. Previously, seven MRMR-selected transcripts were associated with CVDs. Loss of *HBA1* function is linked with coronary artery disease (CAD) [42]. Hypomethylation of *ITGB2* in PBMCs is linked to HF and CAD [43, 44]. SNPs affecting *SELL* are associated with an increased risk of acute coronary syndromes (ACSs) [45]. Blood-based hypermethylation of *ACTB* is associated with the development of coronary heart disease (CHD) [46]. Conversely, hypomethylation increases the risk of stroke [47]. *NCF2* has been used as a diagnostic biomarker for obstructive CAD in PBMCs [48]. Additionally, *NCF2* has been associated with AF [49]. *ICAM3* has been identified as a prognostic biomarker for acute ischemic stroke [50]. *BBLN* has been found DE in damaged hearts [51, 52]. Additionally, we have reported associated CVD risk factors and diseases implicating shared pathways.

**Figure 3.**
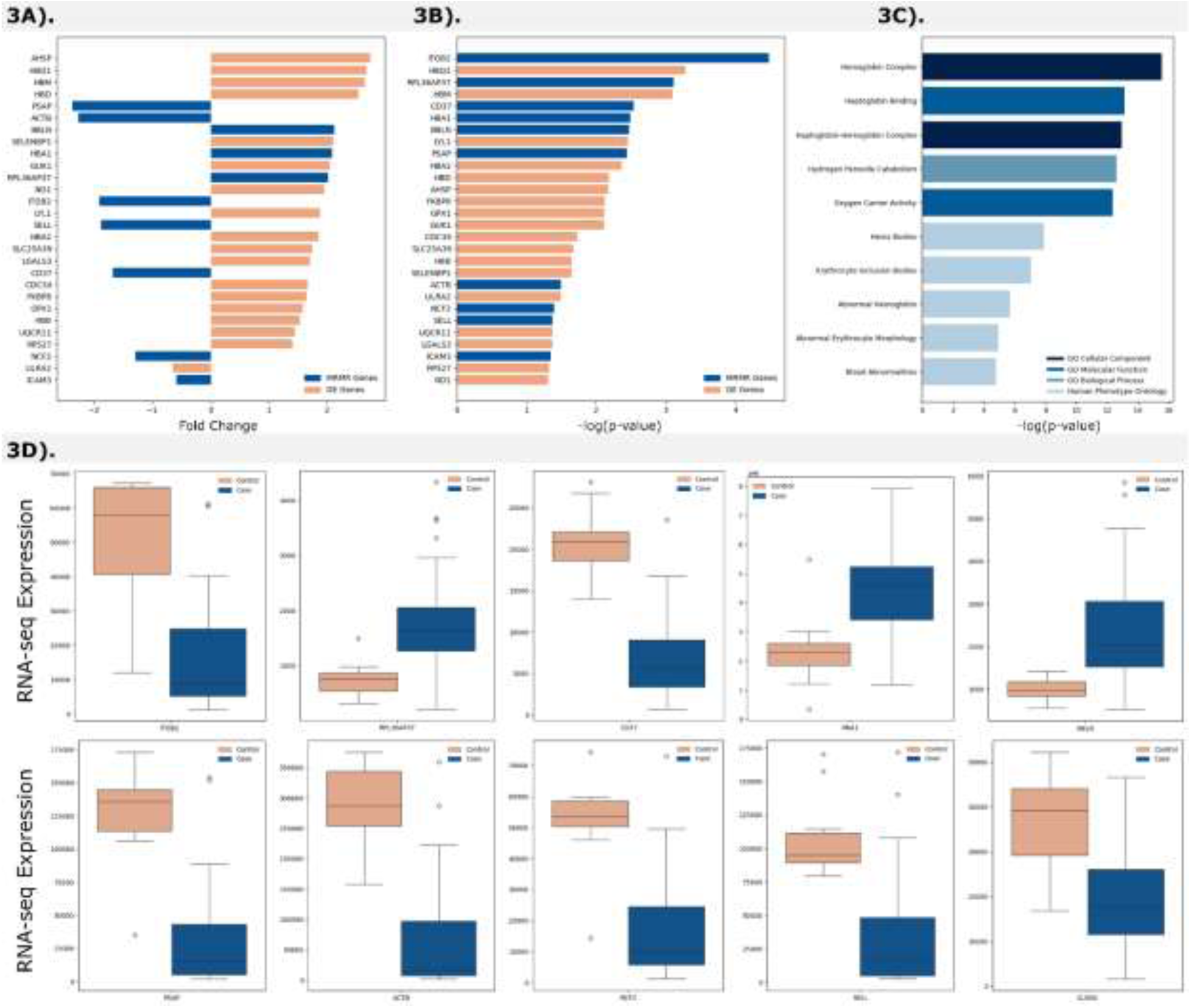
Differentially expressed genes and their expression plots. This figure presents the results of gene expression analysis and that includes, **A)** Fold change in expression level based on differential expression (DE) analysis and redundancy – maximum relevance (MRMR) feature selection; **B)** Significance levels of genes based on DE and MRMR; **C)** Gene annotations for cellular component, molecular function, biological processes, and phenotypic abnormalities; and **D)** RNA-seq expression plots for the ten most significant biomarkers.

We integrated rare, deleterious WGS based SNPs into our patient-specific, ML-efficient profiling. We performed a rigorous search of DEGs and their regulators for SNPs with pathogenic characteristics. Analysis revealed 17 SNPs matching these criteria in our cohort. Figures 4A and 4B demonstrate the CADD Phred scores and gnomAD-sourced allele frequency of these SNPs. Additionally, in Table 2, we report the location, transcripts, and consequences of the 17 SNPs. The distribution of consequences is shown in Figure 4C. The vast majority of our reported SNPs were not reported or scored with uncertain or conflicting pathogenicity in ClinVar. Only *rs115891972* and *rs751011909* were scored benign and likely benign, respectively. Next, using an uncontaminated training dataset, isolated from the testing dataset during feature selection and hyperparameter tuning, we trained three distinct ML classifiers. Our features consisted of 10 MRMR-selected DEGs and 17 SNPs. Our decision tree (DT) classifiers, RF and XGBoost, had perfect abilities to differentiate CVD patients and control individuals. Both classifiers produced 100% and a 1.0 AUC-ROC score. Overall, XGBoost performed the best, considering the Brier score. Our LR model performed worse, scoring 93% accuracy and 1.0 AUC-ROC, but failed to detect the sole healthy individual in our testing dataset. Here, we conclude that DTs are more suitable for single-disease predictions. DTs provide interpretable ML models, capable of handling non-linear relationships and synthesizing various variable types, two strengths necessary for high-dimensional multi-omics datasets. Figures 5A and 5B display our classifiers’ predictions and AUC-ROC. The integrated, multi-omics patient-specific profiles containing SNPs and expression outperformed non-integrated RNA-seq datasets. Previously, we demonstrated 91% accuracy using a comparable dataset with RF and XGBoost classifiers trained on RNA-seq expression [28].

**Figure 4.**
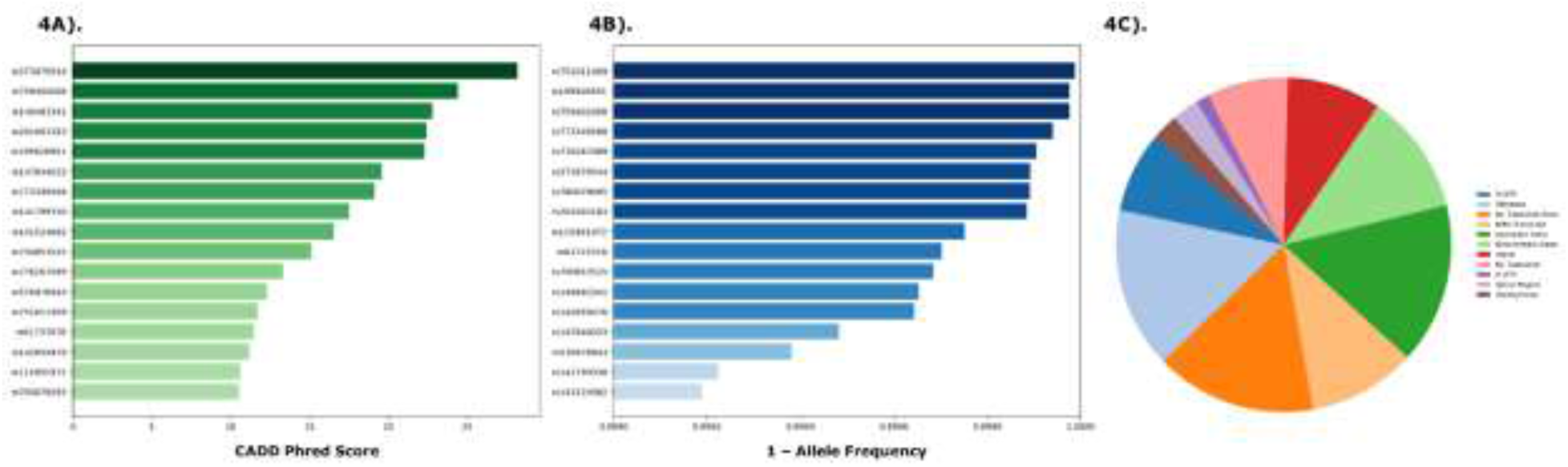
Variant feature selection. This figure presents the rare, deleterious variants affecting our CVD associated biomarkers based on, **A)** Combined Annotation Dependent Depletion (CADD) Score; **B)** Allele frequency obtained from the Genome Aggregation Database (gnomAD); and **C)** annotations of pathogenic single nucleotide variants (SNVs).

**Figure 5.**
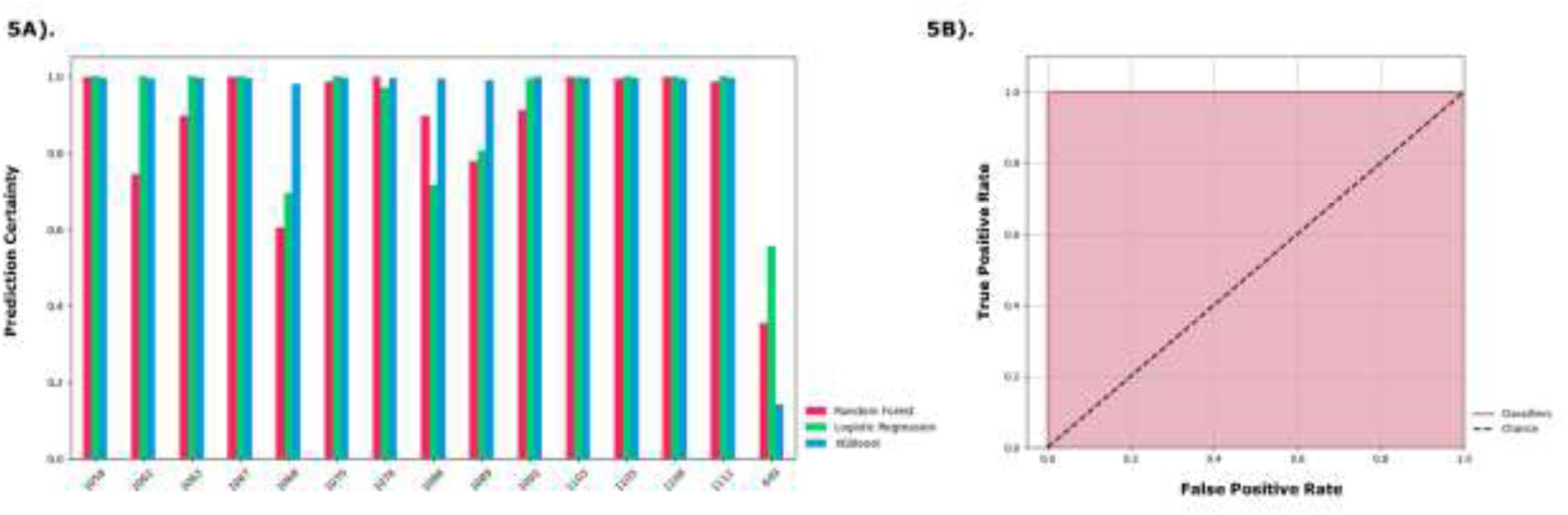
Predictive analysis. This figure presents the predictive confidence of our ML model and that includes, **A)** Predictive certainty of three ML algorithms (Random Forest, Logistic Regression and Xtreme Gradient Boosting) on testing dataset; and **B)** Receiver operating characteristic (ROC) curve denoting the sensitivity and specificity of the classifiers.

**Table 2.** Pathogenic single nucleotide variants (SNPs). This table includes the position of the SNP, its allele, transcripts, consequences (3’ UTR, missense, non-coding (NC) transcript exon, nonsense mediated mRNA decay (NMD) transcript, upstream gene, downstream gene, intron, NC transcript, 5’ UTR, splice region, synonymous), allele frequency, CADD Phred score and corresponding gene.

| SNVs | Position | Allele | Transcript | Consequence | Allele Frequency | CADD Phred | Gene |
| --- | --- | --- | --- | --- | --- | --- | --- |
| rs750853525 | chr21:44891897 | G>A | ENST00000397857<br>ENST00000302347<br>ENST00000475170<br>ENST00000397850<br>ENST00000523323<br>ENST00000355153<br>ENST00000498666<br>ENST00000696946<br>ENST00000397852<br>ENST00000652462<br>ENST00000397854<br>ENST00000479202 | 3'-UTR; Missense;<br>NC Transcript Exon;<br>NMD Transcript;<br>Upstream Gene | 3.152E-04 | 15.13 | ITGB2 |
| rs149483341 | chr21:44900370 | C>T | ENST00000397857<br>ENST00000520389<br>ENST00000523663<br>ENST00000522688<br>ENST00000302347<br>ENST00000523323<br>ENST00000397850<br>ENST00000355153<br>ENST00000498666<br>ENST00000517563<br>ENST00000521987<br>ENST00000397852<br>ENST00000652462<br>ENST00000397854<br>ENST00000522931<br>ENST00000320216 | 3'-UTR; Downstream Gene; Missense;<br>NC Transcript Exon;<br>NMD Transcript | 3.463E-04 | 22.8 | ITGB2 |
| rs141799330 | chr21:44907014 | C>T | ENST00000397857<br>ENST00000521995<br>ENST00000523126<br>ENST00000355153<br>ENST00000498666<br>ENST00000397854<br>ENST00000320216<br>ENST00000520389<br>ENST00000652462<br>ENST00000524251<br>ENST00000479849<br>ENST00000518033<br>ENST00000522688<br>ENST00000523323<br>ENST00000397850<br>ENST00000397846<br>ENST00000521987<br>ENST00000523663 | 3'-UTR; Downstream Gene;<br>Intron; Missense;<br>NC Transcript; NC Transcript Exon;<br>NMD Transcript | 7.756E-04 | 17.52 | ITGB2 |
|  |  |  | ENST00000302347<br>ENST00000517563<br>ENST00000397852<br>ENST00000517819<br>ENST00000522931 |  |  |  |  |
| rs61737078 | chr21:44910770 | G>A | ENST00000397857<br>ENST00000521995<br>ENST00000523126<br>ENST00000355153<br>ENST00000498666<br>ENST00000397854<br>ENST00000320216<br>ENST00000520389<br>ENST00000652462<br>ENST00000524251<br>ENST00000479849<br>ENST00000518033<br>ENST00000522688<br>ENST00000523323<br>ENST00000397850<br>ENST00000397846<br>ENST00000521987<br>ENST00000523663<br>ENST00000302347<br>ENST00000517563<br>ENST00000397852<br>ENST00000517819<br>ENST00000522931 | 5'-UTR; Intron;<br>Missense;<br>NC Transcript; NC<br>Transcript Exon;<br>NMD Transcript;<br>Upstream Gene | 2.975E-04 | 11.49 | ITGB2 |
| rs115891972 | chr14:55137398 | G>A | ENST00000553493<br>ENST00000556263<br>ENST00000254301<br>ENST00000554715<br>ENST00000556438<br>ENST00000553755<br>ENST00000556322 | Intron;<br>NC Transcript; NC<br>Transcript Exon;<br>Splice Region;<br>Upstream Gene | 2.484E-04 | 10.6 | LGALS3 |
| rs759402008 | chr10:71821883 | G>A | ENST00000493143<br>ENST00000495196<br>ENST00000633965<br>ENST00000394936 | Missense;<br>NC Transcript Exon;<br>Upstream Gene | 2.39E-05 | 24.4 | PSAP |
| rs773349568 | chr10:71825851 | T>C | ENST00000493143<br>ENST00000633965<br>ENST00000394936 | Missense; Upstream<br>Gene | 5.97E-05 | 19.12 | PSAP |
| rs776267089 | chr19:13099616 | G>A | ENST00000622520<br>ENST00000585575<br>ENST00000587260<br>ENST00000676441<br>ENST00000360105<br>ENST00000397661<br>ENST00000358552 | Downstream Gene;<br>Synonymous | 9.48E-05 | 13.31 | NFIX,<br>LYL1 |
|  |  |  | ENST00000590120<br>ENST00000590974<br>ENST00000586797<br>ENST00000592199<br>ENST00000588228<br>ENST00000264824<br>ENST00000587760<br>ENST00000693124 |  |  |  |  |
| rs5769789<br>43 | chr19:4933<br>9498 | G>A | ENST00000597602<br>ENST00000598134<br>ENST00000391859<br>ENST00000600121<br>ENST00000595660<br>ENST00000593512<br>ENST00000594743<br>ENST00000597033<br>ENST00000593945<br>ENST00000597852<br>ENST00000358234<br>ENST00000426897<br>ENST00000535669<br>ENST00000602554<br>ENST00000595725<br>ENST00000598095<br>ENST00000377214<br>ENST00000598810<br>ENST00000311227<br>ENST00000539846<br>ENST00000601519<br>ENST00000596426<br>ENST00000323906 | 3'-UTR; Downstream<br>Gene;<br>Intron;<br>NC Transcript;<br>NC Transcript Exon;<br>NMD Transcript;<br>Upstream Gene | 6.18E-04 | 12.28 | TEAD2<br>, CD37 |
| rs1999269<br>51 | chr19:5457<br>4547 | C>T | ENST00000391738<br>ENST00000251377<br>ENST00000391737<br>ENST00000495786<br>ENST00000472992<br>ENST00000251376<br>ENST00000439534 | Missense;<br>NC Transcript Exon;<br>Upstream Gene | 2.39E-05 | 22.3 | LILRA2 |
| rs1476440<br>33 | chr19:5457<br>5483 | T>C | ENST00000391738<br>ENST00000251377<br>ENST00000391737<br>ENST00000495786<br>ENST00000472992<br>ENST00000251376<br>ENST00000439534 | Downstream Gene;<br>Missense; Upstream<br>Gene | 5.173E-04 | 19.6 | LILRA2 |
| rs1415249<br>82 | chr19:5457<br>5519 | G>A | ENST00000391738<br>ENST00000251377<br>ENST00000391737<br>ENST00000495786<br>ENST00000472992<br>ENST00000251376<br>ENST00000439534 | Downstream Gene;<br>Missense; Upstream<br>Gene | 8.107E-04 | 16.54 | LILRA2 |
| rs7600780<br>95 | chr7:55290<br>55 | G>C | ENST00000414620<br>ENST00000432588<br>ENST00000675515<br>ENST00000462494<br>ENST00000473257<br>ENST00000480301<br>ENST00000493945<br>ENST00000464611<br>ENST00000642480<br>ENST00000477812<br>ENST00000674681<br>ENST00000647275<br>ENST00000645576<br>ENST00000676189<br>ENST00000646664<br>ENST00000676319<br>ENST00000645025<br>ENST00000484841<br>ENST00000417101<br>ENST00000425660<br>ENST00000676397<br>ENST00000443528 | Downstream Gene;<br>Intron; Missense;<br>NC Transcript Exon;<br>NMD Transcript;<br>Upstream Gene | 1.084E-04 | 10.53 | ACTB |
| rs1426594<br>76 | chr1:15136<br>5228 | G>A | ENST00000424475<br>ENST00000455397<br>ENST00000447402<br>ENST00000423070<br>ENST00000463664<br>ENST00000427977<br>ENST00000492643<br>ENST00000493560<br>ENST00000465273<br>ENST00000443708<br>ENST00000426705<br>ENST00000470345<br>ENST00000368868<br>ENST00000427867<br>ENST00000455839<br>ENST00000473693<br>ENST00000474352<br>ENST00000458566<br>ENST00000498494 | 3'-UTR; Downstream<br>Gene;<br>NC Transcript Exon;<br>NMD Transcript;<br>Synonymous | 3.56E-04 | 11.2 | SELEN<br>BP1 |
| rs3739795<br>44 | chr1:15136<br>8244 | C>T | ENST00000424475<br>ENST00000455397<br>ENST00000447402<br>ENST00000423070<br>ENST00000463664<br>ENST00000427977<br>ENST00000492643<br>ENST00000493560<br>ENST00000465273<br>ENST00000443708<br>ENST00000426705<br>ENST00000470345<br>ENST00000368868<br>ENST00000427867 | 3'-UTR; Downstream<br>Gene; Missense;<br>NC Transcript Exon;<br>NMD Transcript | 1.074E-04 | 28.2 | SELEN<br>BP1 |
|  |  |  | ENST00000455839<br>ENST00000473693<br>ENST00000474352<br>ENST00000458566<br>ENST00000498494 |  |  |  |  |
| rs751011909 | chr1:183567196 | C>T | ENST00000697330<br>ENST00000420553<br>ENST00000419402<br>ENST00000367536<br>ENST00000697351<br>ENST00000418089<br>ENST00000697329<br>ENST00000413720<br>ENST00000367535<br>ENST00000469280<br>ENST00000495321 | Intron;<br>NC Transcript; Splice Region; Upstream Gene | 1.20E-05 | 11.71 | SMG7, NCF2 |
| rs201003183 | chr1:183586908 | A>T | ENST00000697330<br>ENST00000697558<br>ENST00000367536<br>ENST00000697351<br>ENST00000418089<br>ENST00000697329<br>ENST00000413720<br>ENST00000697353<br>ENST00000367535<br>ENST00000697352<br>ENST00000495321 | Intron; Missense; NC Transcript; NC Transcript Exon; Upstream Gene | 1.153E-04 | 22.4 | SMG7, NCF2 |

At last, we created single-patient profiles of CVD phenotypes using SHAP importances and prediction metrics from our best-performing XGBoost model. Examination of these profiles revealed that the *RPL36AP37* (mean absolute SHAP value = 0.764) and *HBA1* (mean absolute SHAP value = 0.522) demonstrated higher usefulness than other MRMR-selected biomarkers in training the classifier, with BBLN (mean absolute SHAP value = 0.218) the next highest result. *ICAM3* was not utilized by our XGBoost algorithm. Biomarkers contributing more significantly to predictions could indicate disease involvement, leading toward more efficient diagnoses and treatment of CVD.

## 4. Discussion

This study explores the functional impact of AI based multi-omics interactions in CVD. Our DEA disclosed twenty-eight DEGs (Table 1), with twelve out of them associated with a phenotypic variation of CVDs. Further investigation to uncover CVD and non-CVD associations, we found 65% of the total diseases reported for all genes were related to CVDs. Two genes (*HBM* and *GUK1*) had only CVD associations. *HBM* has previously been identified as one of the ten most DEGs for hypertrophic cardiomyopathy (HCM) patients [53]. Additionally, it has been linked to CVD risk factors such as pulmonary arterial hypertension [54] and alpha thalassemia [55]. Upregulation of *GUK1* has been implicated in CHD and HCM [56]. Ten of the twenty-eight DEGs (*HBA1, GPX1, SELENBP1, LGALS3, ND1, ITGB2, ACTB, NCF2, SELL*, and *ICAM3*) were linked to both CVDs and non-CVDs. *HBA1* has been documented to be highly associated with CVDs such as ischemic heart disease [57] and CAD [58]. It has also been linked to other CVD risk factors such as hypertension [59] and alpha thalassemia [59] as well as other non-CVDs such as chronic kidney disease, sickle cell disease [60], and nonalcoholic fatty liver disease [61]. *LGALS3* [62], *ITGB2* [44], and *ICAM3* [50] were all found to be correlated with ischemic stroke and CHD. Upregulation of *GPX1* has been implicated in various complex disorders including but not limited to cardiomyopathy [63], acute myeloid leukemia (AML) [63], and endometrial cancer [64]. Additionally, upregulation of *SELENBP1* has been associated with acute coronary syndrome [65]. It has also been implicated in other phenotypes of CVDs such as myocardial infarction and cardiac arrest [65] as well as non-CVDs including breast cancer and lung adenocarcinoma [66]. Downregulation of *ND1* [67] and upregulation of *ACTB* [46, 68] in the inflammatory pathways are known to be associated with CHD and cardiomyopathy. *ND1* and *ACTB* are also linked to other chronic and heritable diseases such as mitochondrial encephalomyopathy [69] and Parkinson’s disease [70], respectively. Additionally, *NCF2* and *SELL* are reported to be potential diagnostic biomarkers for CAD [71, 72] as well as cancers such as hepatocellular carcinoma [72] and leukemia [73], respectively. While the direct correlation between other complex diseases and CVDs remains unknown, state-of-the-art literature supports the implication of these genes in the inflammatory and immunological pathways shared between these diseases [59, 60, 66, 73]. Future genomic and translational studies are required to understand these relationships.

Sixteen of the twenty-eight DEGs were found to be only associated with non-CVDs based on existing literature (Table 1). Genes such as *HBQ1* [74]*, HBA2* [75]*, CD37* [76], and *LILRA2* [77] are all associated with different types of cancer and other immunological diseases that are documented to have a direct impact on CVD pathophysiology. While these genes are not directly linked to CVDs, further research is required to understand their effects on regulatory elements that might trigger CVDs development. We could not find evidence associating genes such as *BBLN*, *AHSP*, and *HBB* with a CVD or other diseases. However, we reported their implications in CVD risk factors including but not limited to Tetralogy of Fallot [51], beta thalassemia [78, 79], respectively. Other genes such *as RPL36AP37* [80]*, LYL1* [81]*, HBD* [82], *FKBP8* [83], *CDC34* [84], *SLC25A39* [85], *UQCR11* [86], *RPS27* [87], and *PSAP* [88] are all implicated in cancerous and neurological diseases that are not directly associated with CVDs. A detailed list of documented CVD and non-CVD phenotypes associated with our DEGs are available in Table 1.

We validated our findings with peer-reviewed studies (Table 3). Eleven of the twenty-eight genes were found to be upregulated in our DEA as well as existing literature. *HBM* [53], *GUK1* [56], *HBA1* [58], *GPX1* [63], *SELENBP1* [65], and *LGALS3* [62] were all upregulated in different phenotypes associated with CVDs. Other genes such as *HBQ1* [74], *BBLN* [51], *LYL1* [81], *CDC34* [84], and *UQCR11* [86] were implicated in other diseases but their regulation levels also matched existing studies. Additionally, two genes, *SELL* [72] and *CD37* [76] were observed to be downregulated in our DEA and in current studies. Seven of these genes (*HBA1, RPL36AP37, BBLN, ITGB2, ACTB, NCF2, SELL, ICAM3, CD37*, and *PSAP*) were selected using MRMR and later utilized by our AI/ML model to predict CVDs. Upregulation of *HBA1* has been extensively reported to exhibit a strong correlation with ischemic heart disease [57] while loss of *HBA1* function is associated with CAD [55]. In our previous studies, we identified protein coding *HBA1* to be upregulated in CVD patients and significantly expressed in HF patients [14, 25, 26]. Pseudogene *RPL36AP37* assists in regulating DNA replication within eukaryotic cells and in producing ribosomal proteins [89]. Little to no information connecting *RPL36AP37* to CVDs has been reported. However, non-CVD phenotypes associated with altered or loss of *RPL36AP37* function include primary angle closure glaucoma [89], Parkinson’s disease, and certain cancers [80]. Protein-coding *BBLN*, also referred to as bublin coiled-coil protein, serves as a vital regulator of intestinal intermediate filaments, crucial for normal intestinal function, while also playing a role in maintaining cellular organelle architecture and serving as molecular spacers [90]. Induced downregulation of *BBLN* in mice with congenital heart defects leads to further cardiovascular dysfunction and necroptosis via activation of the *CAMK2D* pathway [51]. *ITGB2*, a protein coding gene, encodes a cell-associated signaling molecule particularly involved in leukocyte adhesion and migration of T-cells and neutrophils [91]. *ITGB2* is profoundly expressed within AML patients [43], and hypomethylation of this gene in peripheral blood was linked to HF and CAD [44]. Blood-based hypermethylation of *ACTB*, another protein coding gene, was significantly associated with the development of CHD [46]. While hypomethylation of *ACTB* was found to increase the risk of stroke [92]. Protein coding *NCF2* was found to be significantly upregulated in AF [93] and significantly expressed in CAD patients [48]. *SELL* encodes for selectin, a protein essential for binding and rolling leukocytes on endothelial cells and acts as a primary downstream target for *DYSF*, a protein that, if upregulated, contributes to the pathogenesis of atherosclerotic CVDs [94]. Upregulation of protein coding *ICAM3* is identified as a prognostic biomarker for acute ischemic stroke [95]. *CD37*, protein coding gene, regulates immune response and prevent tumor formation and upregulation in mRNA expression levels is observed in AML patients [76]. *PSAP*, a protein coding gene, plays a significant role in the pathogenesis of atherosclerosis, a key risk factor for CVDs [96]. In particular, elevated *PSAP* expression in plaque macrophages was related to atherosclerosis-linked inflammation in humans [96]. Additionally, these seven genes are linked to other multi-factorial diseases. Further studies are needed to understand non-CVD implication on CVD prognosis and diagnosis.

**Table 3.** Regulation of differentially expressed genes based on literature review. This table includes gene names, their regulation based on the log fold change (LFC), and their regulation based on literature review (up/down). Red-colored text in the regulation based on literature (up/down) refers to cardiovascular diseases while those in blue refer to non-cardiovascular diseases.

| # | Gene | Regulation based on LFC (Up/Down) | Regulation based on literature (Up) | Regulation based on literature (Down) |
| --- | --- | --- | --- | --- |
| 1 | HBM | Up | Hypertrophic cardiomyopathy, alpha thalassemia | N/A |
| 2 | GUK1 | Up | Coronary heart disease, hypertrophic cardiomyopathy | Ischemic stroke |
| 3 | HBA1 | Up | Coronary artery disease, Non-alcoholic fatty liver disease, type 1 diabetes | Acute myeloid leukemia |
| 4 | GPX1 | Up | Cardiomyopathy/cardiac dysfunction, glioblastoma multiforme, renal papillary cell carcinoma, acute myeloid leukemia, ovarian serous cystadenocarcinoma, pancreatic adenocarcinoma, skin melanoma, thyroid cancer, endometrial cancer | N/A |
| 5 | SELENBP1 | Up | Acute coronary syndrome | Renal cell cancer, colorectal cancer, breast cancer, lung adenocarcinomas, prostate cancer |
| 6 | LGALS3 | Up | Heart failure, ischemic stroke, coronary heart disease, atherosclerosis, colorectal cancer, pancreatic cancer | N/A |
| 7 | ND1 | Up | Myeloid leukemia | Maternally inherited coronary heart disease, lactic acidosis, Mitochondrial encephalomyopathy |
| 8 | HBQ1 | Up | Lung adenocarcinoma, secondary progressive multiple sclerosis | N/A |
| 9 | RPL36AP37 | Up | N/A | N/A |
| 10 | BBLN | Up | Tetralogy of Fallot | N/A |
| 11 | HBA2 | Up | Hemoglobin H disease, beta thalassemia | Iron-deficiency anemia, alpha thalassemia |
| 12 | LYL1 | Up | Acute myeloid leukemia, uterine corpus endometrial carcinoma | N/A |
| 13 | HBD | Up | Leprosy | Oral carcinoma |
| 14 | AHSP | Up | N/A | Beta thalassemia, hemolytic anemia |
| 15 | FKBP8 | Up | N/A | N/A |
| 16 | CDC34 | Up | Lung cancer, hepatocellular carcinoma | N/A |
| 17 | SLC25A39 | Up | N/A | N/A |
| 18 | HBB | Up | N/A | Beta thalassemia |
| 19 | UQCR11 | Up | Prostate cancer A, lung adenocarcinoma | Familial hypocholesterinemia |
| 20 | RPS27 | Up | Thyroid cancer | Diamond Blackfan anemia 17 |
| 21 | ITGB2 | Down | Acute myocardial infarction, inflammation and fibroatheroma, atherosclerosis, ovarian cancer, rheumatoid arthritis, osteoarthritis | Coronary heart disease, non-small cell lung cancer, low grade glioma |
| 22 | ACTB | Down | Coronary heart disease, hypertrophic cardiomyopathy | Ischemic stroke, keratoconus |
| 23 | NCF2 | Down | Coronary artery disease, atrial fibrillation, hepatocellular carcinoma, hypertension | Non-small cell lung cancer |
| 24 | SELL | Down | Leukemia | Coronary artery disease |
| 25 | ICAM3 | Down | Ischemic stroke, inflammatory bowel disease | N/A |
| 26 | CD37 | Down | Acute myeloid leukemia, non-alcoholic fatty liver disease | Diffuse large B-cell lymphoma, non-small cell lung cancer |
| 27 | PSAP | Down | Early-onset Parkinson's disease | Atherosclerosis |
| 28 | LILRA2 | Down | Tuberculoid leprosy, rheumatoid arthritis, lepromatous leprosy | N/A |

The integration of multi-omics data, coupled with the multimodal advancements in AI/ML, has the potential to enhance diagnostic and predictive analyses of leading causes of mortality, modifiable risk factors, and other medical insights. State-of-the-art literature have supported the implementation of a genomic language model trained on millions of metagenomic scaffolds to uncover hidden functional and regulatory connections among genes. Additionally, this process also unveils intricate relationships between genes within a genomic region [97]. Another recent study utilized a deep-learning, integrative mass spectrometry framework on metabolomics, with a focus on lipid profiles, to detect lipid content specific to regions and localize lipids to individual cells depends on both cell subpopulations and the anatomical origins of the cells [98]. In epigenetics, recent literature introduced DeepMod2, a comprehensive deep-learning framework designed for methylation detection utilizing the ionic current signal obtained from Nanopore sequencing [99]. It incorporates both a bidirectional long short-term memory model and a transformer model to facilitate rapid and precise detection of DNA methylation from a variety of flow cell types using whole-genome or adaptive sequencing data [99]. Another study showcased a multi-omics analytic platform leveraging genomic, transcriptomic, proteomic, and lipid data to accurately predict adenocarcinoma patient survival [100]. This platform employs an ensemble of algorithms, including support vector machine, random forest, and neural network, to identify disease-associated biomarker panels for downstream predictive analyses [100]. Furthermore, researchers have combined omics data with demographic and clinical information to offer a comprehensive view of cancer prognosis [101]. They created and validated a deep learning framework capable of extracting insights from complex gene and miRNA expression data, enabling accurate prognosis predictions for breast and ovarian cancer patients [101]. All of these approaches offer potential advancements in understanding disease biology and could assist in developing more targeted treatments.

## List of Abbreviations

(ACS): Acute coronary syndrome
(AI): Artificial intelligence
(AF): Atrial fibrillation
(AML): Acute myeloid leukemia
(AUC): Area under the curve
(CVD): Cardiovascular disease
(CAD): Coronary artery disease
(CHD): Coronary heart disease
(CADD): Combined Annotation Dependent Depletion
(CIGT): Clinically integrated genomic and transcriptomic
(DT): Decision tree
(DEGs): Differentially expressed genes
(GO): Gene Ontology
(GSEA): Gene Set Enrichment Analysis
(gnomAD): Genome Aggregation Database
(HF): Heart failure
(HPO): Human Phenotype Ontology
(HCM): Hypertrophic cardiomyopathy
(k-NN): K-nearest neighbors
(LR): Logistic regression
(LFC): Logarithmic fold change
(ML): Machine learning
(MRMR): Minimum redundancy - maximum relevance
(NGS): Next-generation sequencing
(NaN): Not a number
(PBMC): Peripheral mononuclear blood cell
(RMSE): Root mean squared error
(RF): Random forest
(SNPs): Single nucleotide polymorphisms
(SHAP): SHapley Additive exPlanations
(TPM): Transcripts per million
(VEP): Variant Effect Predictor
(WES): Whole exome sequencing
(WGS): Whole genome sequencing
(XGBoost): Xtreme Gradient Boosting

## Acknowledgments

We appreciate great support by Institute for Health, Health Care Policy and Aging Research (IFH), and Robert Wood Johnson Medical School (RWJMS), at Rutgers Health, and Pat and Jim Calhoun Cardiology Center, and Department of Genetics and Genome Sciences, at the UConn School of Medicine, UConn Health. We thank members and collaborators of Ahmed Lab at Rutgers (IFH, RWJMS, RBHS) for their support, participation, and contribution to this study.

We appreciate all colleagues and institutions who provided direct and indirect insight and expertise that greatly assisted the research and development of this project. We acknowledge Rutgers Office of Advanced Research Computing (OARC) for providing access to the Amarel cluster and associated research computing resources.

## Author contributions

Z.A. proposed, led, and supervised this study. Z.A. participated in conceptualization, project administration, funding acquisition, methodology, investigation, resource allocation, data duration, RNA-seq and WGS data processing, quality checking, downstream analysis. W.D. executed formal analysis, and R.N. tested and reproduced results. H.A., E.P., and N.S., participated in research, investigation, and validation of AI/ML results using state of the art literature. S.Z. guided post bioinformatics and AI/ML analysis and evaluated results. B.L. supported overall study including multi-omics data generation. All authors have participated in writing - original draft when Z.A performed review & editing. All authors have approved it for publication.

## Biographical Note

H.A., W.D., E.P., N.S., and R.N. are the Research Assistants/Students at the Ahmed lab, Rutgers IFH/RWJMS.

S.Z. the Assistant Professor at the Department of Biomedical and Health Informatics, UMKC School of Medicine.

B.L. is the Interim Chief Executive Officer (CEO), UConn Health; Executive Vice President for Health Affairs; Dean, UConn School of Medicine; Director, Pat and Jim Calhoun Cardiology Center; and Ray Neag Distinguished Professor of Cardiovascular Biology and Medicine. BL is an internationally recognized cardiovascular physician-scientist and national leader in academic medicine.

Z.A. is the Assistant Professor at the Department of Medicine / Division of Cardiovascular Diseases and Hypertension, Rutgers Robert Wood Johnson Medical School, and Rutgers Health. Z.A. is Core Faculty Member at the Rutgers Institute for Health, Health Care Policy and Aging Research, at Rutgers, The State University of New Jersey. Furthermore, ZA is the Adjunct Assistant Professor at the Department of Genetics and Genome Sciences, School of Medicine, UConn Health, CT.

## Declarations

### Ethical Approval and Consent to participate

Informed consent was obtained from all subjects. All human samples were used in accordance with relevant guidelines and regulations, and all experimental protocols were approved by the Institutional Review Board.

### Consent for publication

Not applicable

### Availability of data and material

All the source code reproducing the experiments of this study are available at GitHub, following web link: <https://github.com/drzeeshanahmed/intelligenes_multi-omics_cvd_analysis>

